# A robust method to isolate *Drosophila* fat body nuclei for transcriptomic analysis

**DOI:** 10.1101/2021.01.27.428429

**Authors:** Vanika Gupta, Brian P. Lazzaro

## Abstract

Gene expression profiles are typically described at the level of the tissue or, often in *Drosophila*, at the level of the whole organism. Collapsing the gene expression of entire tissues into single measures averages over potentially important heterogeneity among the cells that make up that tissue. The advent of single-cell RNA-sequencing technology (sc-RNAseq) allows transcriptomic evaluation of the individual cells that make up a tissue. However, sc-RNAseq requires a high-quality suspension of viable cells or nuclei, and cell dissociation methods that yield healthy cells and nuclei are still lacking for many important tissues. The insect fat body is a polyfunctional tissue responsible for diverse physiological processes and therefore is an important target for sc-RNAseq. The *Drosophila* adult fat body consists of fragile cells that are difficult to dissociate while maintaining cell viability. As an alternative, we developed a method to isolate single fat body nuclei for RNA-seq. Our isolation method is largely free of mitochondrial contamination and yields higher capture of transcripts per nucleus compared to other nuclei preparation methods. Our method works well for single cell nuclei sequencing and potentially can be implemented for bulk RNA-seq.

## INTRODUCTION

The insect fat body is a highly multifunctional tissue that regulates diverse physiological processes such as nutrient storage and metabolic control, immune responses to infection, and production of proteins essential for egg provisioning^1^. This single tissue thus shares function with several vertebrate tissues, including liver and adipose tissue. The fat body is an extremely dynamic tissue that exhibits dramatic expression changes in response to physiological stimulus^2^. It therefore is an important tissue to understand. The diverse functions of the fat body imply that there may be cellular heterogeneity within the tissue, and spatially restricted morphological and functional heterogeneity have previously been observed^3,4^. Single-cell RNA-seq (sc-RNAseq) is a technique that enables transcriptomic profiling of individual cells^5^, which could be invaluable for studying the fat body. However, the success of sc-RNAseq relies on clean, gentle, and rapid dissection of the tissue of interest and dissociation of individual cells. The adult fat body of *Drosophila melanogaster* is a large and fragile tissue that is distributed throughout the body^1^ and is therefore more difficult than other tissues to analyze at a single-cell level. In this manuscript, we compare four methods for isolating fat body cells and nuclei prior to sc-RNAseq. In our hands, isolation of intact fat body cells is infeasibly challenging and results in unacceptably high cellular mortality. Standard protocols to isolate individual nuclei using a sucrose gradient or low-speed centrifugation were successful in capturing nuclei but carried unacceptably high levels of contamination with mitochondria. We ultimately developed a modified method that combines careful tissue dissection and fixation, cell lysis, and nuclear isolation over a sucrose density gradient to generate high-yield, high-purity nuclear isolations that are suitable for transcriptomic profiling.

## RESULTS

### Enzymatic dissociation

Enzymatic dissociation is the most common method to dissociate a tissue into single cell suspension. Several protocols^6^ use papain, collagenase, trypsin or a combination of enzymes to digest a tissue and liberate individual cells. We tested several enzymes including papain, collagenases, trypsin, TrypLE, and Liberase™ in varying enzyme concentration and incubation duration to dissociate the fly fat body tissues into single cells. However, all the methods resulted in a rapid fat body cell death upon incubation with enzymes (Figure 1). Lowering the incubation temperature slows the enzyme activity^7^ and, can lead to lower cell death. However, even when we incubated our tissues at 4°C with Trypsin for 6 hours, we recovered only 15-20% viable cells.

**Figure 1:**
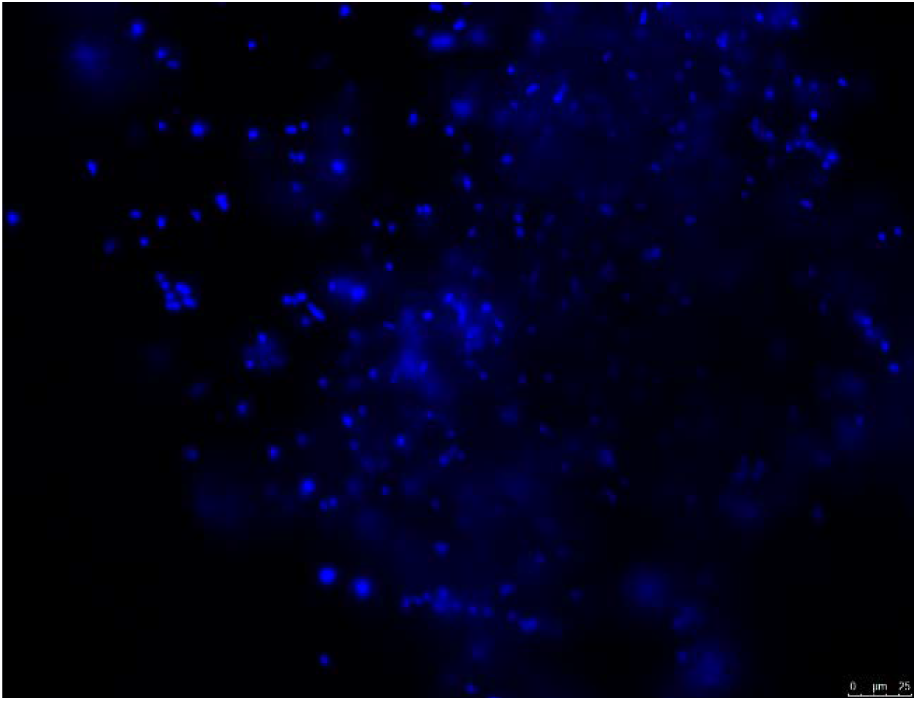
DAPI stained nuclei after enzymatic cell dissociation indicates substantial cell death.

We attempted to use fluorescence-assisted cell sorting (FACS) to recover viable fat body cells after enzymatic dissociation. To label the fat body tissue, we used Gal4-UAS system to drive the expression of EGFP in the adult fat body tissue. We used fat body driver c564-GAL4 (Bloomington Stock Center #6982) with UAS-EGFP to drive the expression in the fat body. After brief enzymatic dissociation (15 minutes) with collagenase I, we used FACS to sort EGFP+ cells from non-fluorescing cells followed by another sort using DAPI live-dead staining to separate live, intact cells from dead cells and debris. We observed variable EGFP expression of EGFP+ cells with no distinct bimodal distribution separating EGFP+ cells from EGFP- cells (Figure 2). Additionally, the sorting revealed large amount of cell debris and DAPI+ nuclei suggesting cell death upon dissociation. These results confirmed that the fat body tissue is extremely fragile, making it almost impossible to use enzymatic dissociation to recover individual viable cells. We also tested the c564-GAL4 driver with UAS-mCherry.NLS (Bloomington Stock Center #38424) as a label to drive fluorescence in fat body nuclei. We found 16N ploidy to be the most abundant nuclei population in the c564-GAL4>UAS-mCherry.NLS compared to 2N and 4N in the control sample (Figure 3). This change in ploidy profile could have an impact on the biology of the tissue rendering this method unfit for our experiments.

**Figure 2:**
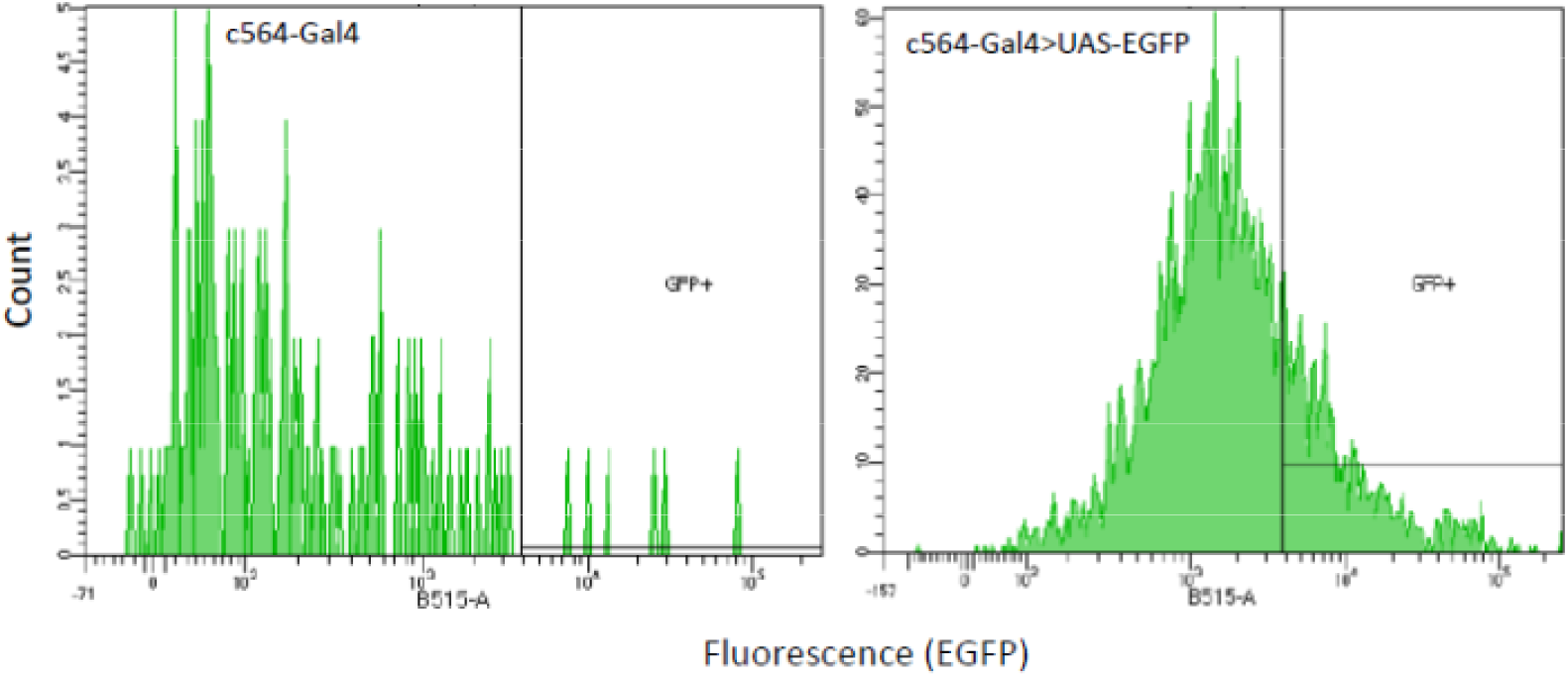
Sorting of EGFP+ cells from EGFP-cells shows variable EGFP expression and lack of separate peak of c564-EGFP+ cells, reflecting few intact viable cells.

**Figure 3:**
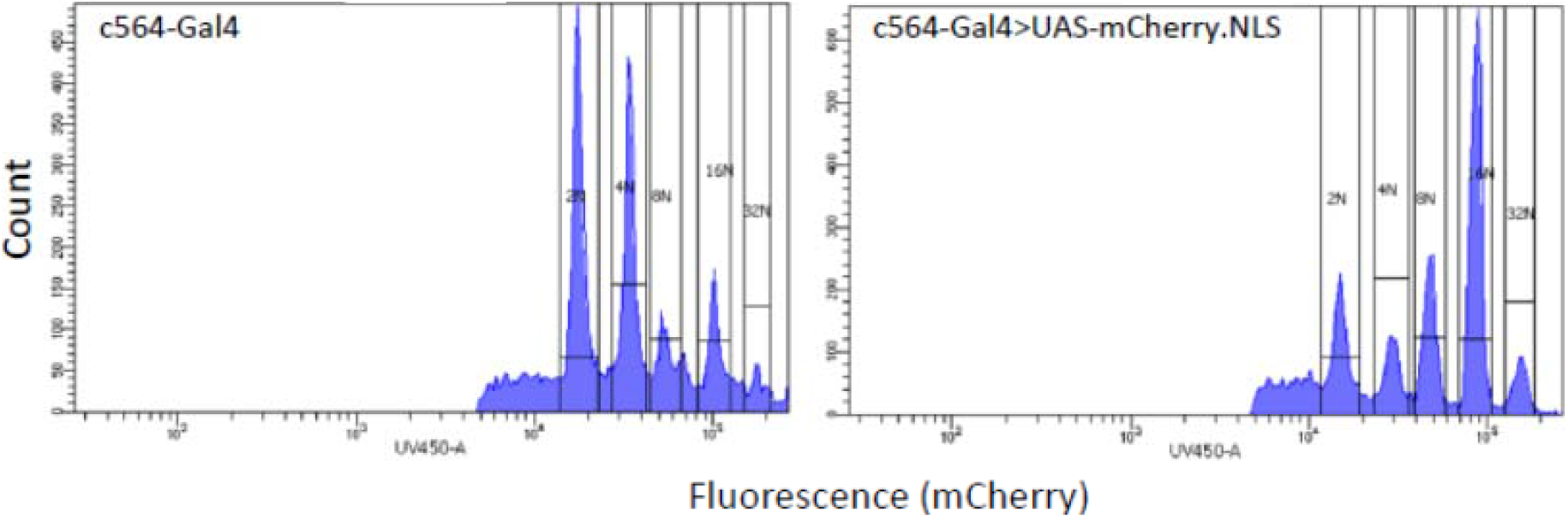
Histogram of DNA content of DAPI+ nuclei show altered ploidy in flies expressing mCherry.NLS+ from the c564 driver.

### Nuclei preparation

Studies^8–10^ have shown that transcriptomic profiles correlate strongly between nuclei and cells, meaning that nascent transcripts in the nucleus are broadly representative of the standing mRNA pool in the cell. Therefore, a nuclear isolation from fat body tissue which could be used for transcriptomic profiling. Our objective was to generate a suspension of nuclei with low contamination from mitochondria and other cellular debris. We tried the following methods:

#### 1. Sucrose cushion gradient centrifugation

In this method, tissue homogenate prepared in hypotonic buffer is passed over a sucrose gradient and centrifuged such that cell debris and other organelles are trapped at specific sucrose densities while the lighter nuclei form a fraction at the bottom of the gradient. We prepared a fat body homogenate using a Dounce homogenizer and hypotonic buffer. The homogenate was centrifuged at 500g for 5 minutes. The pellet was resuspended in PBS containing 2% BSA and the suspension was then passed through a sucrose gradient following the 10X Chromium sucrose cushion protocol^11^ where samples were centrifuged at 13,000g at 4°C. However, our sequencing results showed about 50% mitochondrial reads in our samples, suggesting that the protocol is not ideal for *Drosophila* fat bodies (Figure 4).

**Figure 4:**
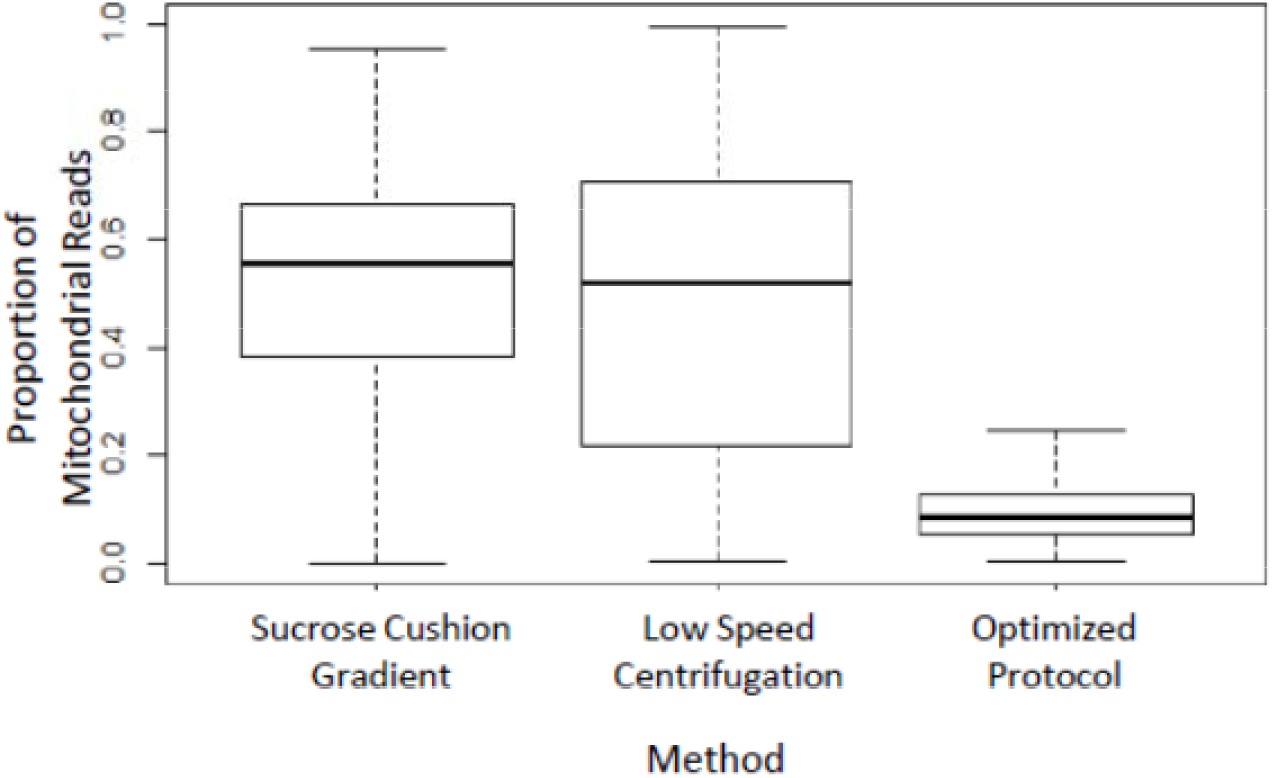
Box plot showing proportion of mitochondrial reads after three different nuclei preparation protocols. Our optimized protocol yields the lowest proportion of mitochondrial reads.

#### 2. Low-speed centrifugation

Low speed centrifugation in hypotonic buffers is typically used^11^ to remove cell debris and mitochondria and pellet nuclei. We centrifuged fly homogenate at 500g for 15 minutes and carefully removed the supernatant. The nuclear pellet was washed three times with hypotonic buffer containing 1U/ul RNase inhibitor and centrifuged at 800g for 15 minutes at 4°C. The pellet was resuspended in PBS containing 2% BSA. Following this protocol, our sequencing results still showed an unacceptably high contamination with mitochondrial reads. Only 25% of the read sets from individual isolated nuclei contained less than 20% mitochondrial reads (Figure 4). The number of median genes obtained per cell was 73 compared to 443 in our optimized protocol described below (Table 1).

**Table 1:** Summary of genes detected in nuclei prepared using three different methods.

|  | Sucrose Cushion Gradient | Low Speed Centrifugation | Optimized Protocol |
| --- | --- | --- | --- |
| Median Genes per Nuclei | 42 | 73 | 443 |
| Total Genes across all Nuclei | 6,923 | 9,339 | 10,125 |

### 3. Optimized nuclei preparation protocol

The inefficiency of pre-existing protocols for the adult fat body prompted us to develop a more effective method to isolate nuclei with less contamination from mitochondria. Using our optimized protocol, we saw dramatic reduction in mitochondrial contamination. Fewer than 5% of the purified nuclei were associated with 20% or higher mitochondrial reads (Figure 4) and there was a considerable improvement in the number of reads and genes obtained per nucleus using the optimized protocol (Table 1). The optimized protocol we developed is as follows:

### Reagents required

#### Adult Hemolymph-like Saline(HLS)

For dissection and storing tissues, hemolymph-like saline of osmolarity suitable for adult flies^12^ was made using the recipe provided by Cold Spring Harbor Laboratories (CSHL, http://cshprotocols.cshlp.org/content/2013/11/pdb.rec079459.full). HLS was filter-sterilized, aliquoted and stored at 4°C until needed.

#### Adult Hemolymph-like Saline (CSHL)

2 mM CaCl_2_
5 mM KCl
5 mM HEPES (pH = 7.4)
8.2 mM MgCl_2_
108 mM NaCl
4 mM NaHCO_3_
1 mM
NaH_2_PO_4_
10 mM Sucrose
5 mM Trehalose
Adjust the pH to 7.5. Filter sterilize and store in aliquots at 4°C

#### Fixation Buffer

4 parts Methanol + 1-part HLS (Pre-chilled at −20°C)

#### Hypotonic Isolation Buffer (HIB)

10 mM HEPES (pH = 7.4)
10 mM KCl
2.5 mM MgCl_2_
0.5 mM Spermidine
0.15 mM Spermine
0.02% Digitonin

#### Hypotonic Sucrose Buffer (HSB)

10 mM HEPES (pH = 7.4)
10 mM KCl
2.5 mM MgCl_2_
0.01% NP40
0.3 M Sucrose

#### Wash and Suspension Buffer (WSB)

1X phosphate-buffered saline (PBS, pH = 7.4)
2% BSA
U/μl RNaseIN

#### Protocol

1. Autoclave micro centrifuge tubes, tips, Dounce homogenizers.
2. Keep micro centrifuge tubes and Dounce homogenizers at 4°C so that everything is chilled before use.
3. Anaesthetize adult flies using CO_2_. Pick a fly using fine forceps and submerge the fly in cold HLS. With the fly submerged in HLS, use the fine forceps and pull at the posterior tip of the abdomen. Use a spring scissors, incise laterally along the cuticle. Open the cuticle and carefully remove the ovaries, gut, and Malpighian tubules, exposing the fat body layer attached to the cuticle.
4. Transfer the dissected fat body tissue, still attached to the cuticle, to Fixation Buffer. Incubate on dry ice for 5-10′.
5. Transfer the dissected tissue to 1 mL Dounce homogenizer containing 1mL HLS on ice.
6. Repeat steps 3-5 to pool desired number of dissected fat bodies. We pooled 40 tissues for the data shown above. *Note*: 40 abdomens reliably yield approximately 10^6^ nuclei as counted by DAPI staining.
7. Carefully remove the HLS from the tube leaving enough liquid such that tissues do not dry out.
8. Add 1 mL of ice-cold HIB. Allow tissues to swell up in hypotonic buffer for 5 minutes.
9. Dounce the tissues gently twice with loose pestle and once with tight pestle. Keep the pestle submerged in the buffer during lysis avoiding bubble formation.
10. Transfer tissues along with hypotonic isolation buffer to 7 mL Dounce homogenizer with a wide-mouth P-1000 pipet tip. Wash the 1 mL homogenizer with 1mL of HIB and transfer to 7 mL homogenizer. Add another 2 mL of HIB and Dounce twice with tight pestle.
11. Transfer lysate to 5 mL centrifuge tube. Pass lysate through a 22G syringe 3-5 times carefully. Do not allow the abdominal cuticle to pass through the syringe.
12. Transfer supernatant to 2 mL LoBind^®^ centrifuge tubes (Eppendorf #022431048) and spin at 800g for 7′ @ 4°C.
13. Carefully remove and discard the supernatant without disturbing the pellet.
14. Gently resuspend pellet in HSB and centrifuge at 1500g for 10′ @ 4°C.
15. Carefully remove supernatant and resuspend the pellet in 500 μl of WSB.

We further removed cellular debris using sucrose gradient centrifugation. Using Sigma kit (NUC201-KT), we followed the protocol using instructions provided by 10X Chromium^11^. The protocol is as follows:

16. Prepare Sucrose Cushion Buffer I (SCB I) by mixing 2.7 ml Nuclei PURE 2M sucrose cushion solution (Component NUC201-KT) and 300 μl Nuclei PURE sucrose cushion buffer (Component NUC201-KT). Keep it on ice.
17. Transfer 500 ul of SCB I to a 2.0 ml LoBind^®^ microcentrifuge tube. Keep it on ice.
18. Take 900 ul of SCB I and add it to 500 μl of nuclei suspension from Step 15 in the main protocol. Mix thoroughly but gently.
19. Carefully overlay the nuclear suspension mixed with SCB I (from Step 18) onto the 500 μl SCB I in the LoBind^®^ microcentrifuge tube (from Step 17). Do not mix the two layers.
20. Without disturbing the layers, transfer the LoBind^®^ tube to a microcentrifuge pre-chilled at 4°C.
21. Centrifuge samples at 3,500g for 20′ at 4°C.
22. Carefully remove the tube from the centrifuge and place on ice.
23. Remove 1950 ul of the supernatant, leaving 50 μl in the tube. If the nuclei pellet is not visible then leave around 100-150 μl of supernatant.
24. Resuspend the pellet in WSB to a total volume of 500 μl.
25. Pass the sample through a Flowmi^®^ cell strainer (BAH136800040) with 40 μM pore size. The sample may need to be passed twice if there is excess debris.
26. Immediately proceed to the sequencing protocol or any other downstream use of the purified nuclei.

## DISCUSSION

Here we provide an optimized method for preparing a nuclear suspension suitable for transcriptomics. Our method uses a combination of detergents and gentle centrifugations over a sucrose gradient to generate a purified nuclear suspension with low mitochondrial contamination. Fixing the samples immediately upon dissection preserves the quality of the RNA and results in excellent transcriptomic data.

Single-cell sequencing technology requires dissociation protocols that can provide high yield of intact living cells from which a standing mRNA pool can be reliably recovered. However, cell dissociation protocols are delicate and the precise protocol needs to be specifically optimized for the tissue type of interest. This may prove challenging for fragile tissues such as the insect adult fat body, whose cells which are unstable and rapidly die upon dissociation. On the contrary, nuclei can be isolated from any cell or tissue type. Since the nascent transcript pool in the nucleus is strongly correlated with the standing mRNA pool in the cytoplasm^8–10^, sequencing nuclei can provide a reliable alternate strategy for sequencing any tissue at single cell resolution.

Although our protocol was specifically developed for *Drosophila melanogaster*, we expect that it can be broadly applied. The fat body is a major tissue in all insects and performs several key functions including immune response, metabolism, production of egg yolk and vitellogenin, and xenobiotic detoxification. Single-cell/nucleus sequencing provides a tremendous opportunity to study this crucial insect tissue and understand the cellular basis for functional diversity within the tissue. We expect that that the protocol we present here can be readily applied to the fat body of non-*Drosophila* insects and can be adapted for other tissue types as well.

## Acknowledgments

Dr Christopher Donahue, manager of the flow cytometry facility in the College of Veterinary Medicine, Cornell University provided technical support with cells/nuclei sorting.

